# Survival time and prognostic factors in dogs diagnosed with haemangiosarcoma in UK first opinion practice

**DOI:** 10.1101/2024.12.07.627327

**Authors:** C. Taylor, G.J. Barry, D.G. O’Neill, A. Guillén, P. Pickard, J. Labadie, D.C. Brodbelt

## Abstract

Visceral haemangiosarcoma is considered clinically aggressive in dogs, with perceived poor prognosis often leading to euthanasia at presentation. This study aimed to determine survival times and prognostic factors in dogs with haemangiosarcoma under first-opinion care.

Dogs diagnosed with haemangiosarcoma in first-opinion practice in 2019 were identified in VetCompass electronic health records (EHRs) and manually examined to capture variables potentially associated with survival. Median survival time (MST) from diagnosis was calculated for the whole population and based on the primary tumour location. Binary logistic regression was used to explore differences between dogs that died on the day of diagnosis and those that survived ≥1 day. Cox proportional hazards modelling explored factors associated with time to death in dogs that survived ≥1 day.

Across all haemangiosarcoma cases (n=788), overall MST was 9.0 days (95%CI:5.0-15.0, range: 0-1789) and proportional 1-year survival was 12.0% (95%CI:9.7-15.0%). Dogs with splenic (MST=4.0 days, 95%CI 0.0-9.0) and cutaneous haemangiosarcoma (MST=119.0 days,95%CI:85.0-248.0) had MST greater than 0 days. Male sex and increasing tumour size were associated with increased hazard of death while cutaneous location and surgery were associated with reduced hazard of death.

A very short survival time was identified for haemangiosarcoma under first-opinion care. Although survival time was longest for cutaneous cases, the actualised prognosis was poor overall for haemangiosarcoma. This study provides valuable information for contextualised care and dialogues with clients in first-opinion practice.

## Introduction

Canine haemangiosarcoma is a malignant neoplasia of vascular endothelial cells or bone marrow-derived-progenitor cells and is most commonly found affecting spleen, heart, liver, cutaneous and subcutaneous tissues (1). Incidence estimates vary substantially, depending on the population studied, but two UK studies, one of dogs under first-opinion care and one of insured dogs, have estimated 0.007-0.25% of dogs were affected (2,3). Confirmatory diagnosis of haemangiosarcoma is achieved through histopathological diagnosis on biopsies (1). As it is a highly aggressive neoplasm, metastases are common at presentation. Staging assesses the number of tumours, lymph node involvement, distant metastaes and whether the tumour has ruptured to determine if a haemangiosarcoma is Stage I, II or III (1,4,5).

Treatment options include mainly surgical excision of the mass and/or chemotherapy (1,4). Use of radiotherapy has also been reported for cardiac and cutaneous haemangiosarcomas (1,5). Whilst surgical excision can be curative for cutaneous forms, other haemangiosarcomas have have a poor prognosis (1,6). Factors associated with survival outcomes can be broadly split into non-modifiable (such as animal signalment, tumour stage, tumour size and tumour location) and modifiable factors (such as medical or surgical treatments undertaken) (Table 1). The direction and magnitude of these associations can vary by haemangiosarcoma location (7–9).

**Table 1.** Published factors reported to affect survival post haemangiosarcoma diagnosis ↑ = increased survival time, ↓ = reduced survival time and - = no effect.

| Modifiable effect | Effect/s on survival | References | Unmodifiable effect | Effect/s on survival | References |
| --- | --- | --- | --- | --- | --- |
| Cutaneous-complete tumour resection | ↑ | Shiu et al., (2011); De Nardi et al., (2023) | Cutaneous-Tumour size >4cm | ↓ | De Nardi et al., (2023) |
| Cutaneous – incomplete resection + radiotherapy | ↑ |  | Cutaneous – tumour in sun exposed location/ solar induced form | ↑ | Szivek et al., (2012) |
| Cutaneous- pre-surgical resection chemotherapy (of non-removable mass) | ↑ | Wiley et al., (2010) | Cutaneous – concomitant neoplasia | ↑ | Nobrega et al., (2019) |
| Cardiac – doxorubicin | ↑ | Mullin et al., (2016) | Cutaneous – breeds predisposed | ↑ | Nobrega et al., (2019) |
| Cardiac – surgery | ↑ | Yamamoto et al., (2013) | Cutaneous – female sex | ↓ | Nobrega et al., (2019) |
| Cardiac- surgery and chemotherapy | ↑ | Yamamoto et al., (2013) | Splenic-thrombocytopaenia at diagnosis | ↓ | Masyr et al., (2021) |
| Splenic – chemotherapy after splenectomy | ↑<br>Or<br>- | Wendelburg et al., (2012)<br>Moore et al., (2017) | All – disease stage I<br>All – disease stage II or III | ↑<br>↓ | Ward et al., (1994);<br>Wendelburg et al., (2015); |
|  |  |  |  |  | Masyr et al., (2021);<br>De Nardi et al., (2023) |
| Splenic-metronomic chemotherapy post splenectomy | ↑<br>Or<br>– | Batschinsk i et al., (2018)<br>Alexander et al., (2019) | All – presence of metastases at diagnosis | ↓ | De Nardi et al., (2023) |
| Splenic-polysaccharopeptide | ↑ | Cimino Brown et al, (2012) |  |  |  |
|  | – | Gedney et al., (2022) |  |  |  |

Survival time for these patients vary, according to tumour location and treatment pursued. Cutaneous locations are reported to have the longest survival times (between 172 to 1189 days) and cardiac haemangiosarcoma the shortest (between 7 to 189 days) (Mullin et al., 2016). The reported median survival time for splenic haemangiosarcoma ranges from 23 to 259 days (1,7,10).

Clinical staging is reportedly the most predictive factor associated with survival time in cutaneous and splenic haemangiosarcoma (1,7,11,12).

In visceral forms of haemangiosarcoma, metastases have frequently occurred at point of diagnosis (1). However, for cutaneous masses, metastases are less common and surgical resection can be curative (1,13,14). For both visceral and cutaneous forms, dogs with a Stage I haemangiosarcoma have increased survival time when compared to Stage II or III tumours and Stage II tumours have increased survival versus Stage III (1,7,11,12).

For cutaneous masses, smaller tumour size has been associated with increased survival time (1,14), however this association has not been seen for splenic haemangiosarcoma (7,15). Size does not appear to have been explored as a prognostic factor for cardiac or hepatic haemangiosarcoma locations.

Whilst certain breed types (such as Retrievers and Shepherds) are at increased risk of diagnosis with haemangiosarcoma (1,16,17), less is known about the effect of breed on survival time. For cutaneous haemangiosarcoma, breeds that are predisposed to developing haemangiosarcoma (such as Whippets, Bull Terriers and Dalmatians) have increased survival time when compared to breeds that are less commonly diagnosed with cutaneous haemangiosarcoma (1,13,18). The short coat on the predisposed breed types is associated with developing solar induced cutaneous haemangiosarcomas which are associated with increased survival time (1). For splenic haemangiosarcoma, no survival difference was seen when large and small breed dogs were compared (19) and further breed associations do not appear to have been explored.

For visceral haemangiosarcomas there are no reported associations between sex and neuter status and survival time. One cutaneous haemangiosarcoma study found females to have reduced overall survival time when compared to males (13).

In splenic haemangiosarcoma, chemotherapy and method of chemotherapy administration post-splenectomy had mixed effects on survival time, with some studies reporting an increased survival time associated with chemotherapy (19,20) while others did not (7,8). For cardiac cases, surgical management and/or chemotherapy are reportedly infrequently pursued due to poor prognosis and the high skill level required for cardiac surgery (9). However, small studies that pursued surgery and/or chemotherapy did report increased survival time (9,21). Alternative therapeutics such as *Yunnan baiyao* and *Coriolus versicolor* have been explored but evidence for their effect on survival time has not been reported (1,22,23).

Previous studies examining haemangiosarcoma survival time and factors affecting survival time in dogs have largely been limited to smaller numbers of dogs, referral centre populations or submissions to tumour registries (11,14,18,21,24). Consequently, findings from these studies may generalise poorly to dogs under first opinion care. Therefore, this study aimed to utilise the VetCompass database of first opinion practice attending dogs to explore survival time of haemangiosarcoma (overall and by location) and factors associated with survival time. It was hypothesised that cutaneous haemangiosarcomas have the longest survival time of all locations and that increasing tumour size is associated with reduced survival time. Additionally, it was hypothesised that the subset of dogs that died on day of diagnosis were significantly different from dogs with ≥1 day of survival and therefore there would be factors associated with dying on the day of diagnosis.

## Methods

### Study population and case identification

The study population included all dogs under first opinion care at participating VetCompass clinics in 2019. Dogs were defined as under veterinary care if they had at least one electronic health record (EHR) —such as a clinical note, treatment, or bodyweight measurement—recorded during 2019 or in both 2018 and 2020. Available data fields include species, breed, date of birth, sex, and neuter status, clinic ID, corporate veterinary group, as well as date-specific information on free-text clinical notes, bodyweight, and billed item treatments.

Potential haemangiosarcoma cases (candidate cases) were identified using the following search terms in the clinical notes: (haemangios*, hemangios*, HSA, angiosa*, haemangiosarcoma∼1, hemangiosarcoma∼1, hameangios*). A haemangiosarcoma case was defined as a dog with evidence of a final clinical diagnosis (with or without histopathological confirmation) of haemangiosarcoma in the EHRs. Only incident cases (those diagnosed between January 1 2019 and December 31 2019) were retained for this study. Ethical approval for the VetCompass programme was granted by the Social Science Research Ethical Review Board at the Royal Veterinary College (SR2018-1652).

### Clinical record coding

A total of 4,997 candidate cases were randomly ordered and each EHR was manually reviewed to determine if it matched the haemangiosarcoma case definition. Candidate cases were excluded if haemangiosarcoma was listed only as a differential diagnosis or if there was no final diagnosis. Confirmed haemangiosarcoma cases had additional clinical information extracted and coded. Information from the coded variables was then grouped into further categories for case description, survival time calculations and regression modelling (Table 2).Fixed values of breed, sex, neuter status, date of birth and clinic and corporate group IDs were downloaded automatically from the EHR. The information coded and its groupings are shown in Table 2. Cases were restricted to those first diagnosed at ≥5 years old, as haemangiosarcomas (amongst other cancers) have been shown to be rare in young dogs (3,25).

**Table 2.** Information extracted from clinical records of haemangiosarcoma cases diagnosed in 2019 in VetCompass EPRs. The ‘Variable’ column shows what information was captured from EPRs and what it was transformed or grouped into for regression analyses (‘Grouped variables’).

| Variable | Grouped variables |
| --- | --- |
| Age | Age at first diagnosis (continuous),<br>Age at first diagnosis |
| **Neutering** | Neutering status when diagnosed<br>Time between neutering and diagnosis |
| **Breed** | - Top 20 most common breed types in VetCompass in 2019<br>- Breeds with $\geq 5$ cases<br>- Breeds with $\geq 10$ cases<br>- Ancestral breed grouping (26)<br>- Genotype breed grouping (27) |
| **Clinical signs** | Clinical signs (CS):<br>o Haematological<br>o Cardiac<br>o Respiratory<br>o Gastrointestinal<br>o Urinary<br>o Mass specific<br>o Other specific<br>o Non specific<br>o No CS reported<br>o Effusion present |
| **Diagnostic tests performed** | - Diagnostic test groups:<br>o Imaging<br>o Laboratory tests<br>o Sampling<br>o Cardiac diagnostics<br>o Abdominal diagnostics<br>o No diagnostics done |
| **Surgical management options** | - Did they have any surgery performed?<br>- Did they have surgery and medical management performed? |
| **Medical management options** | - Did they have any medical management?<br>- Type of medical management:<br>o Cardiac<br>o Alternative supplementation<br>o Transfusion<br>o Haemostatic<br>o Palliative |
| **Tumour specifics** | - Any tumour location:<br>o Splenic<br>o Hepatic<br>o Cardiac<br>o Cutaneous<br>- Presence of metastases?<br>- Metastatic tumour location:<br>o Abdominal metastases<br>o Thoracic metastases<br>o Cranial metastases<br>o Soft tissue metastases<br>o Lymphatic metastases<br>o No location specified<br>- Main interest:<br>o Cardiac<br>o Splenic<br>o Hepatic<br>Tumour size<br>Maximum tumour size (quartiles, terciles and continuous) |
| **Referral for advanced clinical management** | Did the dog visit a referral centre/specialist? |
| **Death** | Survival time (days)<br>Mechanism of death not taken forward into analyses |
| **Indices of Multiple Deprivation (IMD)** | IMD quintiles |
| **Urban- rural classification** | Urban- rural classification |
| **Clinic ID** | Clinic ID |

Due to the aggressive and invasive nature of visceral haemangiosarcoma it was considered unlikely to be able to establish whether a tumour location was primary or metastatic mass based on the EHR information so tumours were additionally grouped more broadly into whether there was any tumour present at the spleen, heart, liver or cutaneous tissues (‘any tumour location’ variables in tumour specifics row in Table 2).

### Survival time

Survival was calculated from date of first diagnosis to death or until the end of the study or the animals last available EHR (if the last EHR was before December 31, 2023). Animals that had not died were censored for survival analysis on December 31, 2023. Median survival time (MST, days) and 1-year survival rates (range, % and 95% confidence intervals, CI) were calculated separately for: all haemangiosarcoma cases and all cases that lived ≥1 day after first diagnosis. These calculations were also performed separately by tumour location for splenic, hepatic, cardiac and cutaneous tumours. Kaplan–Meier survival (*i.e*. time-to-event) curves were constructed to describe time to death from haemangiosarcoma diagnosis for all cases and then separately for each tumour location group.

### Factors associated with dying on day of diagnosis

Binary logistic regression modelling was used to ascertain whether there were distinct associations between a range of factors and an outcome variable of either dying on the day of presentation and or dying ≥1 day after presentation.

Initial univariable binary logistic regression modelling assessed for potentially significantly associated variables. Explanatory variables with liberal associations with survival beyond day of diagnosis (p<0.2) were carried forward for multivariable binary logistic regression modelling (28). A correlation matrix was built and Variance Inflation Factor (VIF) was used to assess for collinearity between potentially significant explanatory variables. Substantial collinearity was considered present with correlation ≥0.7, VIF> 5 (28).. For collinear variables, each of the variables were assessed individually in the model and the variable with smallest model AIC value was retained for further modelling. The multivariable model was built using a manual stepwise forwards construction, approach where variables with the smallest p-value from the likelihood ratio test (LRT) and/or smallest Akaike Information Criteria (AIC) value were included first. Variables were retained in the model if the model was deemed improved with variable inclusion through a significant likelihood ratio test (LRT) result. AIC values were used to compare non-nested models that were built with different collinear variables to determine which model balanced goodness of fit and complexity best. Pairwise interactions were assessed for variables in the final model and retained if significant (LRT p <0.05). Performance of the final model was assessed through Area Under the Curve (AUC) and model fit to data was assessed by a Hosmer-Lemeshow test, where p>0.05 indicates that the model adequately fits the data (28). A random effect of clinic ID was added to the model. Intraclass coefficient (ICC) was calculated to compare level of variance within a clinic to level of variance between clinics. Statistical significance was set at the 5% level (28)

### Cox regression modelling

All dogs with ≥1 day survival after diagnosis (n=407) were considered in Cox proportional hazards regression modelling to evaluate the hazard of death from diagnosis with haemangiosarcoma. Explanatory variables with liberal univariable association with hazard of death with haemangiosarcoma (P<0.2) were carried forward for multivariable Cox proportional hazards regression modelling (28). A correlation matrix for potentially significant explanatory variables, as described for the binary logistic regression model approach. Clinical signs were excluded from multivariable Cox regression modelling due to their lack of independence. Both age at diagnosis (years) and maximum tumour size (cm) recorded were both normally distributed and appeared to increase linearly with hazard of death and both variables were explored as a continuous variable and as categorical variables by quartiles. The categorical maximum tumour size variable was taken forward into multivariable Cox regression modelling due to the large number of missing values of tumour size (n=258) whereas age was retained as a continuous variable as there were no missing records for this.

Multivariable model building used a manual stepwise forwards construction approach. Variables were retained in the model if the model was deemed improved with variable inclusion through a significant likelihood ratio test (LRT) result and the smallest AIC value for separate models built with collinear variables. Pairwise interactions between final model explanatory variables were evaluated and retained if their inclusion significantly improved the model (LRT p value <0.05). The proportional hazards assumption (*i.e*. the assumption that the hazard ratios (HRs) for category variables remain constant over time) was tested through assessment of Schoenfeld residuals and visualisation of survival curves in Kaplan-Meier plots for each variable (28). Schoenfeld residuals were assessed for individual variables in the final model and for the global model. If the global model did not violate the proportional hazards assumption (p>0.05) then this model was used. Model performance and fit to the dataset was assessed through Concordance Index and Xu and O’Quigley R^2^ values (28–30). Statistical significance was set at the 5% level (28).

All data cleaning and analyses was performed in R Studio (R Core Team, Vienna, Austria) using the following packages: survminer, dynpred, ggsurvfit, ggplot2, finalfit and coxr2.

## Results

### Study population

From the study population of 2.2 million dogs, there were 4,997 candidate haemangiosarcoma cases. Of these, 788 cases met the case definition for newly diagnosed haemangiosarcoma in 2019. Once age restricted to dogs over five years, the median age at first diagnosis was 10.6 years (intraquaritle range, IQR, 9.1-12.1 years). Nearly three-quarters of the population were neutered (n=577,73.4%). Male dogs comprised 53.5% of the cases (n=422) and females 45.6% (n=359). The three most common breeds were: Labrador Retriever (96/788, 12.2%), German Shepherd Dog (n=84, 10.66%) and Staffordshire Bull Terrier (n=43, 5.5%). Overall, crossbred dogs were the most common dog type (n=177, 22.5%).

There were 29 tumour locations recorded with the most common ones being spleen (n=489, 62%) and liver (n=198, 25.1%) followed by cutaenous haemangiosarcoma (n=146,18.5%).

Most EHRs reported at least one clinical sign at presentation, with 28 (3.5%) dogs having no clinical signs recorded at presentation. A wide range of clinical signs were recorded, with 64 distinct clinical signs noted. Most commonly clinical signs were non-specific: lethargy (n=309, 38.6%), anorexia/hyporexia (n=287,35.8%) and abdominal distension (n=197, 25.0%) (Figure 1).

**Figure 1.**
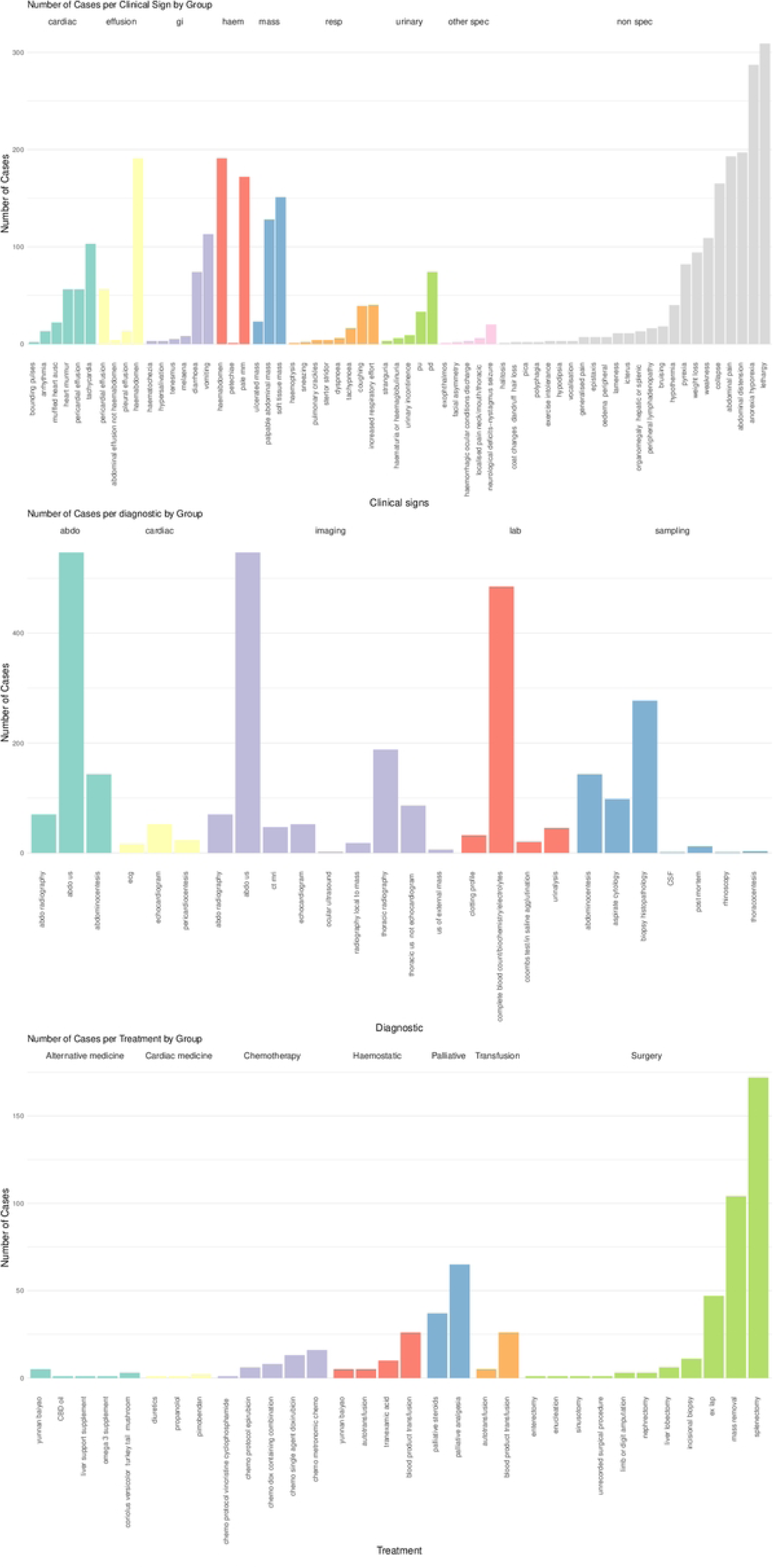
Grouping of clinical variables coded from electronic health records of dogs presenting with haemangiosarcoma to VetCompass clinics in 2019 (n=788). Bar chart colours indicate the group that an individual variable was grouped into: Clinical signs grouped according to their main body system affected (top bar plot), diagnostic tests performed (middle bar plot) and treatment options undertaken (bottom bar plot).

The most commonly performed diagnostic tests were: abdominal ultrasound (n=547, 68.3%) and blood tests (n=485, 60.5%). Effusion analysis was performed in 166 patients (20.7%), of these the majority were abdominocentesis (n=143, 17.8%) and less commonly pericardial effusions (n=23,2.9%). In one-third of patients, histopathological diagnosis was performed (n=277, 34.6%).

For staging purposes, 30% (n=238) of patients had their tumour size recorded, only 21.4% (n=169) of patients had presence of metastases recorded and of these dogs, only 7 (0.9%) recorded metastases to lymph nodes. Therefore, tumour staging was not taken further in the survival analyses. Median tumour size was 6.0cm (IQR=3.5-10 cm, range 0.2-23.0 cm).

Surgical excision was the most commonly performed treatment (n=276/788, 35.0%), with splenectomy being the most common type of surgery (n=172/276, 62.3%). Palliative care (analgesia or steroid administration) was the next most common treatment (n=102, 12.9%). Chemotherapy (either high-dose or metronomic) was uncommon (n=42, 5.3%). Of those, anthracycline-based(doxorubicin or epirubicin) protocols were most commonly used (27 dogs, 64% of chemotherapy dogs) followed by metronomic cyclophosphamide (16 dogs, 38% of chemotherapy patients).

Whether death was the result of euthanasia or unassisted was recorded in most patients(n=676, 85.8%), with euthanasia being the mechanism of death in most cases (n=624, 92.3%).

### Survival time

Survival time across all haemangiosarcoma cases ranged from 0-1789 days (IQR 0-74.25 days). The median survival time (MST) of all cases was 9 days from diagnosis (95% CI 5-15 days), with a 1-year survival rate of 12% (9.7-15%) (Table 3). Dogs with cardiac and hepatic haemangiosarcoma had the lowest MST at 0 days (95%CI 0-0) for both (Figure 2). The 1-year survival rate for these locations was 3% (95% CI 0.8-12%) and 3.9% (95% CI 1.8-84%) respectively. The MST for splenic haemangiosarcoma was 4 days (95% CI 0-9 days). Cutaneous haemangiosarcoma had the longest MST at 119 days (95% CI 85-248 days), with a 1-year survival rate of 35% (95% CI 28-44%).

**Figure 2.**
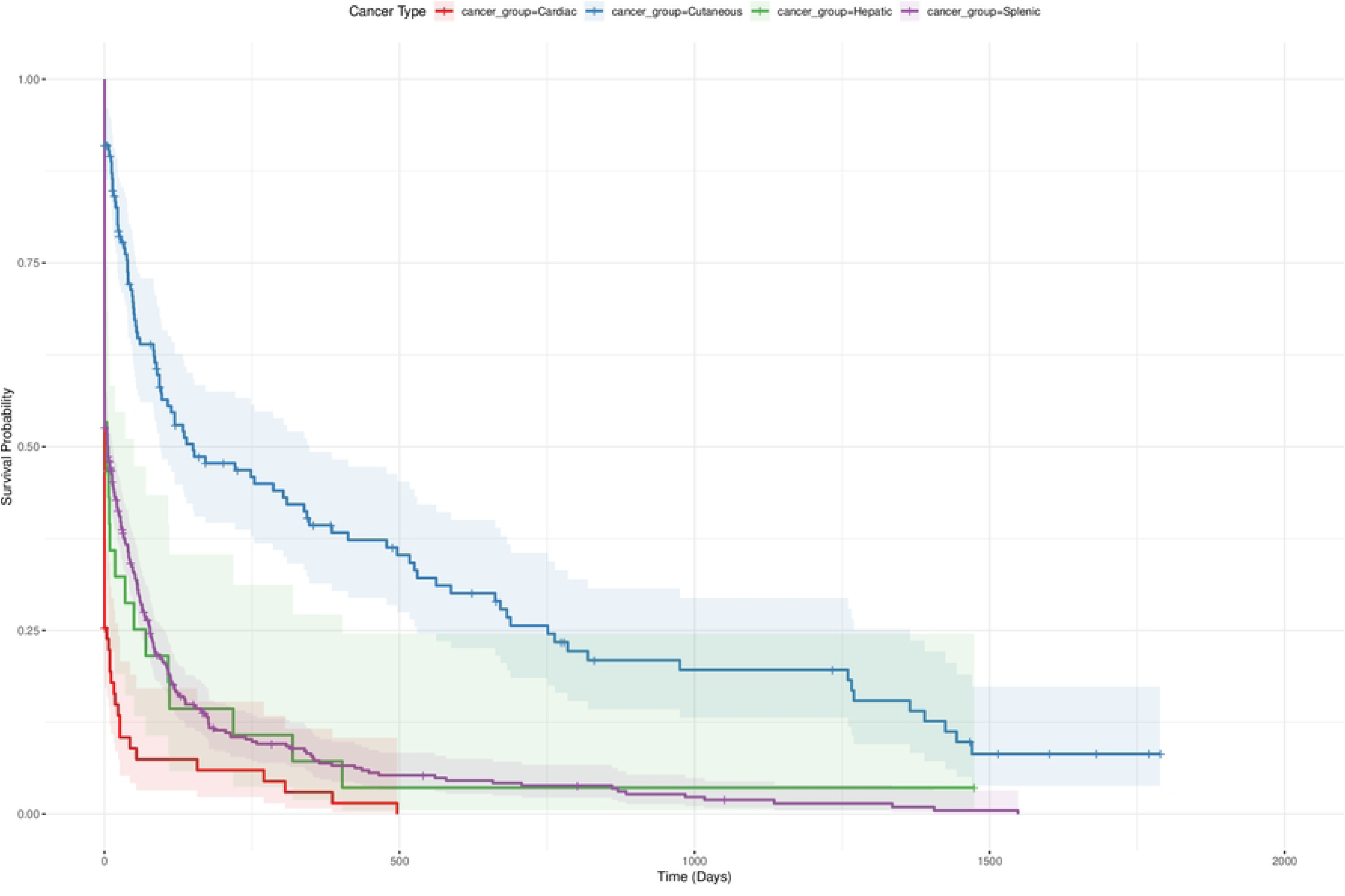
Kaplan-Meier survival curve estimating the survival time of haemangiosarcoma cases diagnosed in first-opinion practices in VetCompass in 2019 separated by location affected: cardiac, cutaneous, hepatic, splenic. Day 0 designates day of first diagnosis. Shaded areas indicate the 95% CI for each curve and crosses indicate censored cases.

**Table 3.**
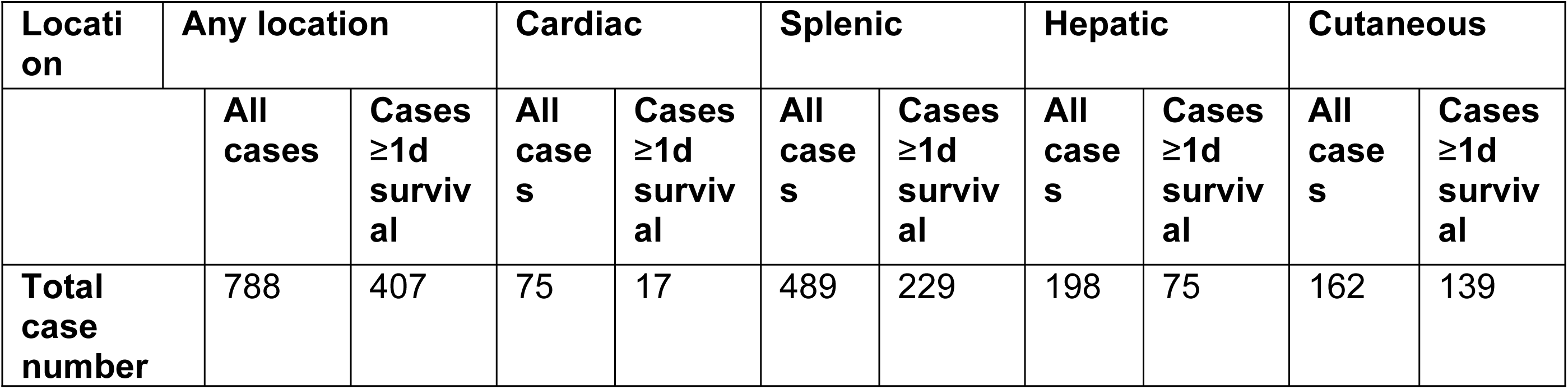
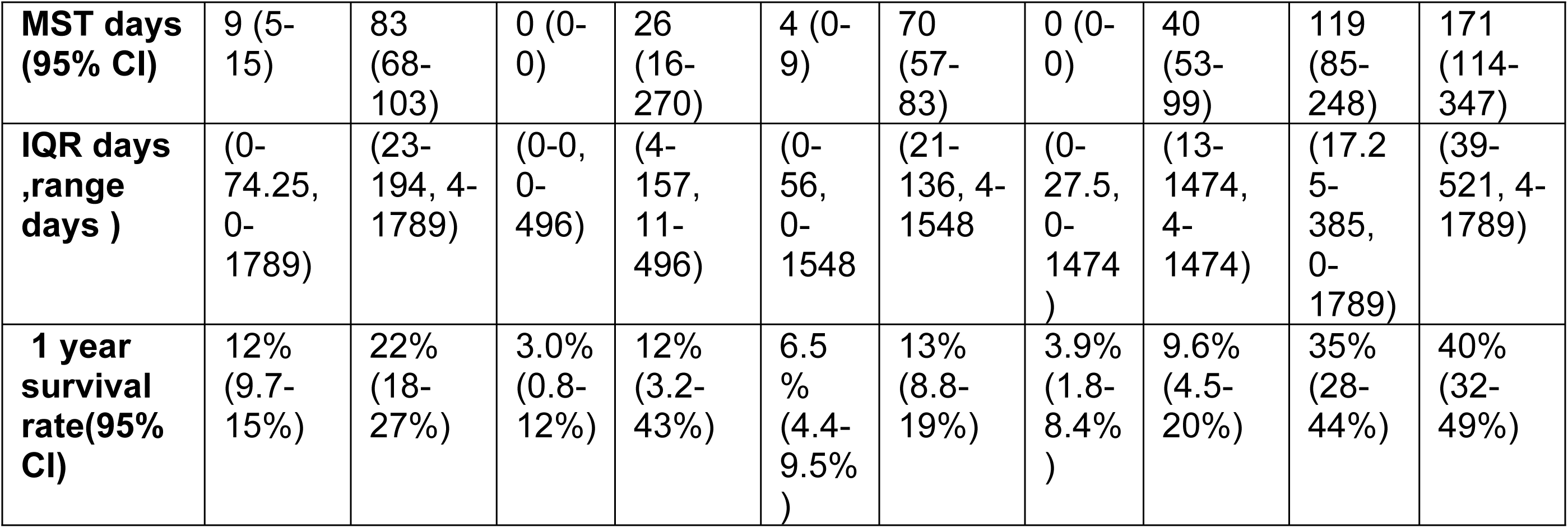
Summary table of case numbers, median survival time (MST), interquartile range (IQR) and 1-year survival rate for haemangiosarcoma cases diagnosed in first opinion practices in VetCompass in 2019. Cases are grouped by location: any, cardiac, splenic, hepatic and cutaneous. Cases are divided further to separate all cases and cases with ≥1 day of survival time post-diagnosis.

### Factors associated with death on day of diagnosis

Due to the high proportion of dogs not surviving for ≥1 day after diagnosis (n=381/788, 48.3%) identified in survival time calculations it was hypothesised that there were two distinct populations of haemangiosarcoma cases. Therefore, binary logistic regression was performed to ascertain risk factor differences between dogs with <1 day of survival (n=381) and those with ≥1 day of survival after diagnosis (n=407). In univariable logistic regression comparing these two groups, 37 variables were found to be potentially significant (LRT p-value<0.2) that were taken forward to multiple logistic regression (Table A1 in Supplementary Material). Of these variables, there were 52 pairs of correlated variables which were assessed in non-nested models to decide on which of the pair to include. The final model comparing differences between the dogs with <1 day survival and ≥1 day survival retained 8 variables: if the patient had surgery or medical treatment, haemangiosarcoma location, whether they visited a referral centre, whether there were mass associated clinical signs present), whether there were haematological abnormality clinical signs present, whether abdominal diagnostic tests were performed and whether there were any clinical signs recorded at presentation. Additionally, corporate group was forced into the model to account for the structure of the data and Clinic ID was retained as a fixed effect (Figure 3). The model showed no evidence of poor fit to the data (Hosmer-Lemeshow p>0.05) and good discrimination (AUC =0.93). There was a small amount of clustering at the clinic level (ICC=0.11).

**Figure 3.**
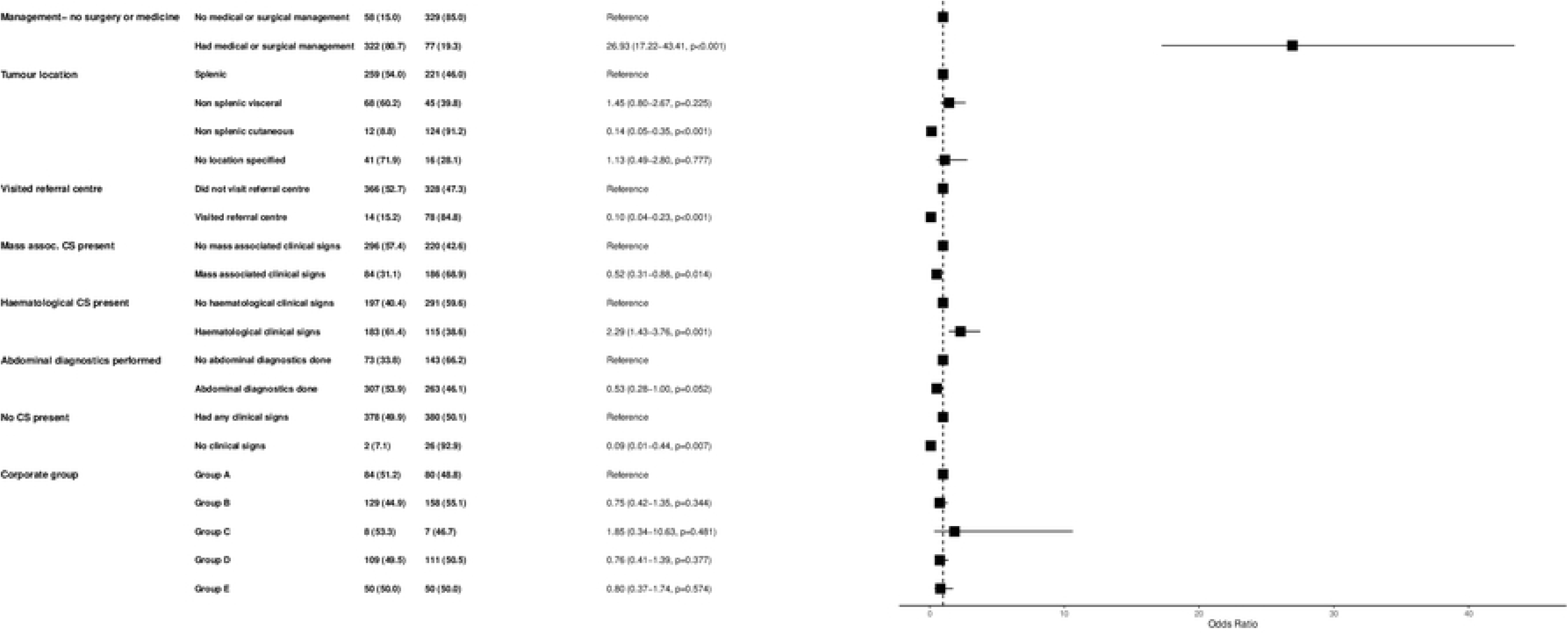
Forest plot of final multivariable logistic regression model of haemangiosarcoma cases diagnosed in first-opinion practices in VetCompass in 2019 who died on day of diagnosis (‘Day 0’, n=381) and those who died ≥1 day of survival post diagnosis (‘Day ≥1’, n=407). Day 0 and Day≥1 columns represent number of animals and percentages for each category in the model. OR indicates the odds ratio for each category and its 95% confidence interval (CI) and p-value.

Dogs that received any surgical or medical management of their haemangiosarcoma were more than 25 times more likely to die on day of diagnosis than those that did not (OR=26.93, 95% CI 17.22-43.41). The presence of haematologically abnormal clinical signs at presentation was associated double the likelihood of dying on day of diagnosis (OR=2.29, 95% CI 1.43-3.76). Dogs with no clinical signs recorded at presentation were ten times less likely to die on day of diagnosis (OR=0.09, 95% CI 0.01-0.44). Dogs with mass associated clinical signs at presentation were half less likely to die on day of diagnosis (OR=0.52, 95% CI 0.31-0.88).

Dogs who visited a referral centre as part of their treatment were more than 10 times less likely to die on day of diagnosis (OR=0.10, 95% CI 0.04-0.23). A cutaneous location was associated with a seven-fold reduction in odds of dying on day of diagnosis compared to a splenic haemangiosarcoma (OR=0.14, 95%CI 0.03-0.23). Dogs with any abdominal diagnostic tests performed were half as likely to die on day of diagnosis (OR=0.53, 95%CI 0.28-1.00). None of the corporate group variables were significant.

These large variations in odds ratios between dogs that died on day of diagnosis and dogs that survived for ≥1 day indicated that these two dog groups were distinct and not suitable to be combined for survival analyses. In order to evaluate factors associated with survival with haemangiosarcoma diagnosis beyond first day of diagnosis, only dogs with ≥1 day of survival time were taken forwards.

### Hazard for death with haemangiosarcoma in dogs that survive for ≥1 day after diagnosis

Of the 63 variables examined in univariable analyses, 41 were liberally significant (LRT p-value <0.2). Variables describing clinical signs at presentation were not taken forward due to their lack of independence as factors, leaving 43 factors to be assessed in multivariable modelling (Table A3 in Supplementary material). Of these, 28 variables were potentially correlated which were assessed in non-nested models for inclusion.

In the final Cox proportional hazards regression model, five variables were retained: whether a patient had any form of surgery, location of haemangiosarcoma, sex, maximum tumour size and the corporate group the clinic belonged to. Corporate group not significant but was forced into the model to better account for the structure of the data. Additionally, a frailty term of clinic ID was incorporated to further account for data structure. The overall model and individual independent variables were assessed for violation of proportional hazards. All individual variables and the overall model satisfied this assumption. The final model performed well, with a pseudo-R^2^ of 0.26 and a Concordance Index of 67.

Hazard of death increased in as the maximum tumour size increased, with greatest hazard in dogs with haemangiosarcomas between 8-23 cm when compared to those with 0.2-4.3cm tumours (HR=1.94 95% CI 1.19-3.18). Male dogs had an increased hazard of death (HR=1.32, 95%CI 1.05-1.65) compared to female dogs. Dogs with any surgery performed had more than half the reduction in hazard of death when compared to those that didn’t (HR=0.48, 95%CI 0.38-0.62). When compared to cardiac haemangiosarcoma cases, dogs with a haemangiosarcoma at other locations had reduced hazard of death with haemangiosarcoma, although this was only significant for cutaneous. Dogs with a cutaneous haemangiosarcoma had half the hazard of dogs with cardiac (HR=0.53, 95% CI 0.40-0.69) (Figure 4).

**Figure 4.**
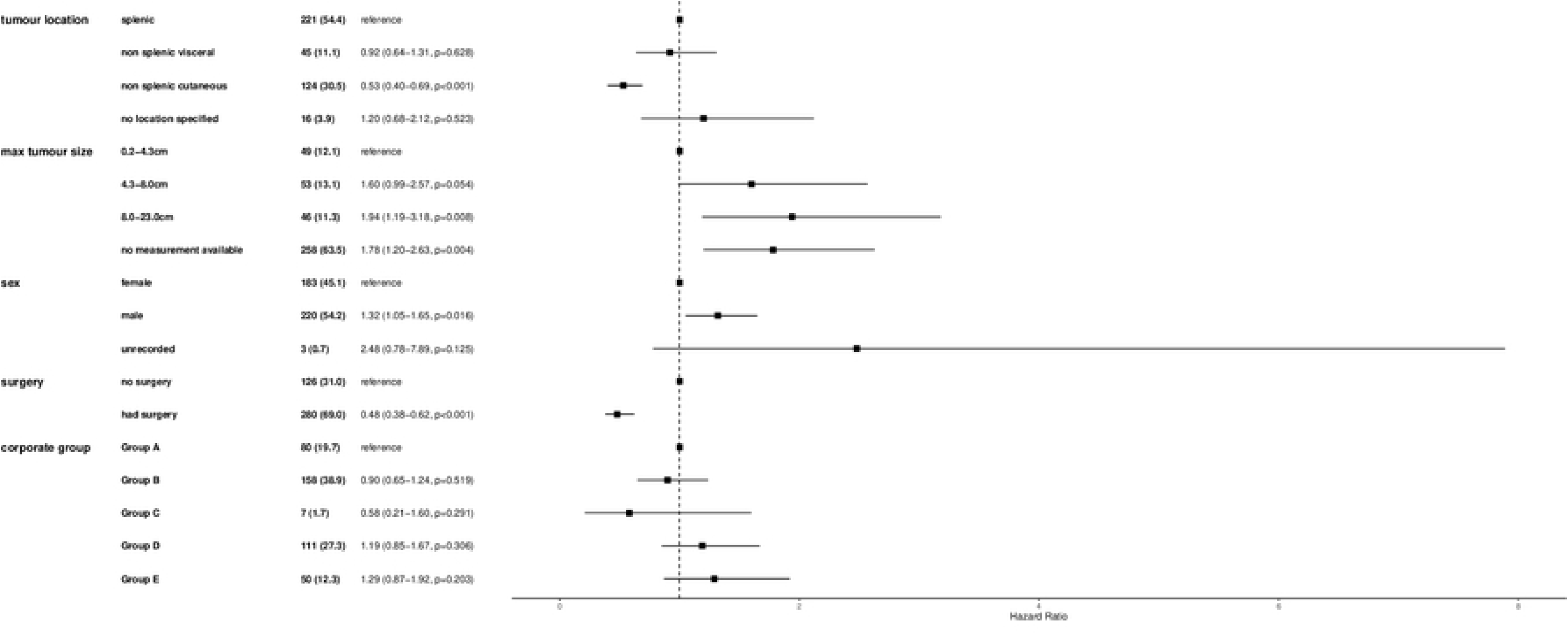
Forest plot of final mixed effects multivariable Cox proportional hazards regression results for factors associated with time to death with haemangiosarcoma in cases diagnosed in first-opinion practices in VetCompass in 2019 with ≥1 day of survival post diagnosis (n=407). Cases (%) column indicates the number of cases for each category in the model. HR indicates the hazard ratio for each category and its 95% confidence interval (CI) and p-value. The final model included a frailty term of Clinic ID.

## Discussion

This is the first study exploring haemangiosarcoma survival time and factors associated with death in a first opinion practice attending in the UK. Existing studies have focussed on referral care populations which exhibit biases(1,7,13,15), as such this work aimed to explore haemangiosarcoma survival from a first opinion perspective. This study identified a short MST overall for haemangiosarcoma (nine days) and for each main tumour location. In hazard regression modelling of dogs with ≥1 day of survival after diagnosis, dogs with haemangiosarcoma of a cutaneous location and dogs that had any surgical management performed had half the hazard of death. Male sex and increasing tumour size were associated with increased hazard of death. Despite not being able to fully disentangle clinical presentation and other influences on haemangiosarcoma, the findings provided here can assist general practitioners to provide location specific and UK first opinion specific survival time estimates.

### Survival time after diagnosis with haemangiosarcoma

Visceral haemangiosarcomas are commonly regarded as aggressive neoplasms with a poor prognosis (1,4). Short median survival times (MST) were consistently seen across all haemangiosarcoma locations in this study, When compared to existing studies, each haemangiosarcoma location had a substantially shorter median survival time in this study. Whilst dogs with cutaneous haemangiosarcoma had the longest survival time of all locations here (MST=119 days) it was still shorter than previous studies, which reported MSTs between 172 to 1189 days (12–14,24). Likewise, in the literature cardiac and splenic cases still have very short MST (7-189 days and 30-259 days, respectively) but this was substantially longer than what was seen here (0 days and 4 days, respectively) (7,10,21,31,32).

In most previous studies exploring haemangiosarcoma survival, studies include small case numbers primarily treated in specialist centres and presence of referral bias is likely (7,8,14,31). This could have affected survival time through the availability of more advanced therapies, owner more motivated to pursue intensive treatment options and less financial concerns if attending a referral centre. Additionally, patients attending a referral centre might have had to present to their primary veterinarian clinically stable enough for referral. Furthermore, some studies excluded patients that did not meet a minimum survival length or did not survive to discharge which would have skewed for a longer MST (7,15). When MSTs were calculated for cases with ≥1 day of survival after diagnosis in this Vet Compass population, MSTs were similar to those reported in previous studies.

The potential reasons for short survival times are multifactorial. Haemangiosarcomas are reported to be generally aggressive tumours, with high metastatic rates (4). In the study population here, a low proportion of dogs were receiving chemotherapy and many presented with severe clinical signs leading to potential inferences about low quality of life in haemangiosarcoma patients. However, the common prevailing view of extremely poor prognosis for haemangiosarcoma could be promoting frequent euthanasia at presentation and therefore leading to a self-fulfilling prophesy and low survival times. From reading clinical records it is difficult to be certain on the nuances of why euthanasia has been performed, with factors such as perceived prognosis, financial considerations and emotional distress being more difficult to capture than patient clinical presentation. As a result, it is not clear the proportional contribution of effects from these non-clinical factors on a decision to euthanise are challenging to derive and therefore the impact on survival times reported here.

For cutaneous haemangiosarcoma, the MST reported in the current results was at the lower end of previously reported MSTs in the literature (12–14,24,33). This may have reflected that our categorisation of cutaneous haemangiosarcomas encompassed a wider range of non-visceral, more aggressive forms such as subcutaneous and intramuscular haemangiosarcomas. Additionally, solar radiation induced forms of haemangiosarcoma have been associated with increased survival time than non-solar induced (1,18). Whilst the solar induced aspect of cutaneous haemangiosarcoma biology was not assessed for in cases here, due to reduced sunlight hours and intensity and UV levels solar-induced haemangiosarcoma cases may be less frequent in northern Europe, meaning survival time may be reduced in cutaneous haemangiosarcomas in this current UK study (34,35).

In previous referral centre studies tumour staging has been shown to be a key prognostic factor (1). However, tumour staging in accordance with adapted WHO guidance (tumour size, evidence of regional and distant metastases) was rarely recorded in clinical records in this study. Therefore, it was not possible to explore how the MSTs reported here relate to cancer stage

### Factors associated with dying on day of diagnosis

The binary logistic regression model aimed to explore clinico-pathologic and treatment factors associated with dogs that died on day of diagnosis versus dogs that survived for at least one day. Two factors were associated with dying on day of diagnosis: having surgical or medical management of their haemangiosarcoma and the presence of haematologically abnormal clinical signs. The management of haemangiosarcoma being associated with increased odds of death on day of diagnosis seems counter-intuitive initially, however this most likely reflects unstable patients requiring urgent surgery (for example, an actively bleeding splenic mass requiring emergency surgery) being associated with poorer outcomes. Likewise, the reduced likelihood of visiting a specialist centre in dogs that die on day of diagnosis also supports this, as dogs may need to be clinically stable before travelling for specialist care. As anticipated, based on survival times in existing literature (1,13,14), presence of a cutaneous haemangiosarcoma was associated with reduced odds of dying on day of diagnosis.

This model indicated that dogs dying on day of diagnosis had different associations with both modifiable and unmodifiable haemangiosarcoma factors than those that survived longer.

### Factors associated with time to death with haemangiosarcoma

Previous survival analyses on haemangiosarcoma tended to focus on single locations for haemangiosarcoma. However, factors identified in this current study represent associations across all types of haemangiosarcoma. Cutaneous haemangiosarcomas were at significantly reduced hazard of death when compared to splenic haemangiosarcoma cases. The association of cutaneous haemangiosarcoma location with lowest hazard of death was consistent with median survival times reported in the literature across haemangiosarcoma locations (1,7,8,14,15), but this has not been explored through multi-location haemangiosarcoma survival analyses with a single study before.

In the current study, increasing tumour size was associated with increased hazard of death. Whilst larger masses (>6cm) have been shown to be associated with a reduced survival time in cutaneous haemangiosarcoma (14), in several previous splenic haemangiosarcoma studies size has not been associated with prognosis (7,15). One reason for this lack of association in previous studies was survival to discharge post-splenectomy as an inclusion criterion might select for smaller masses as these might make better surgical candidates. Additionally, tumour size may only be associated with prognosis for certain haemangiosarcoma locations. Since other studies exploring tumour size have focused on a single haemangiosarcoma location, our model which includes all locations makes interpreting this size association more complicated. For example, cardiac haemangiosarcomas have a very short survival time but are limited by cardiac anatomy to the extent they can grow to (31).

Male dogs had an increased hazard of death with haemangiosarcoma in the current study. An association between sex and haemangiosarcoma survival time or diagnosis does not appear to have been previously reported (31,36). Besides cancers of reproductive organs with their obvious sex associations there does not appear to be strong evidence for sex being associated with canine cancer survival time. This contrasts with humans, where men are often reported at increased risk of diagnosis and also poorer survival times across several cancers (37–39). In human oncology, the exact mechanism of sex on both diagnosis and survival is not well understood (40). There are sex hormone associations but also lifestyle and environmental differences between human sexes which contribute to this variation (37–39). Sex hormones are involved in the regulation of physiological process such as stem cell renewal and the immune response and androgen receptor expression has been identified in angiosarcomas (a human neoplasm similar to haemangiosarcoma in dogs), indicating a potential sex hormone association for this cancer (40). However, the extent to which the sex difference is translatable to veterinary is unclear given how commonplace neutering is, which minimises the potential role of sex hormones (41). Neuter status was not retained in the final hazards model, further indicating that there are other factors beyond sex hormones contributing to this finding.

No significant effect on survival time was seen in the current study when medical treatments such as chemotherapy or alternative medicines (such as *Yunnan baiyao* or *Coriolus versicolor*) were utilised. The use of these therapeutics was uncommon (40 dogs for chemotherapy, 9 for alternative medicine), likely leading to a lack of power to detect an effect. However, as there is currently minimal evidence to suggest alternative medicines improve survival time it is also probable this association would not exist in the data here (23,36).

Previous studies reported some breed associations with haemangiosarcoma diagnosis but there is limited work evaluating the breed associations on survival after diagnosis (17,42). Only cutaneous haemangiosarcomas has been previously reported to have some breed associations with survival time (13). However, this was not found to be significant whether a dog survived ≥1 day or not, nor with the survival time overall, in agreement with existing literature (14,19).

### Limitations

Whilst working with large volumes of clinical records in VetCompass has the advantage of providing high number of cases there are some limitations to working with anonymised, retrospective records. For example, full laboratory records such as histopathology on masses or blood tests are often not available in the EPRs. In addition, the current first opinion study included many patients that did not have a definitive diagnosis pursued (only one-third had histopathology performed) or blood samples taken (only 60% of cases). This meant that the study was unable to assess the effect of variables associated with prognosis identified in other studies such as the mitotic count of a mass, Ki67 expression or thrombocytopaenia (8,11).

Relying on clinical diagnoses rather than histopathological diagnosis could have lead to inclusion of misdiagnosed patients, as not all bleeding spleens are malignant masses and not all malignancies are haemangiosarcoma. Application of the ‘double two-thirds’ rule (where two-thirds of splenic masses are malignant and of these a further two-thirds are haemangiosarcoma) here indicates that the potential for misdiagnosis due to a lack of definitive diagnosis could have had an impact on survival time and factors associated with it identified here (10). Meta-analyses by Schick and Grimes (2022) indicated that both splenic malignancies and haemangiosarcoma might be more common than the rule suggests (43).

Achieving a definitive diagnosis can prognosticate better for owners and avoid a potentially unwarranted euthanasia (if benign). However, as this is often not feasible in acute, critical clinical presentation scenarios and in first opinion practice, presumptive diagnoses were included in the current analyses. Additionally, cytological diagnosis has limitations (such as blood contamination of samples) and an invasive biopsy may be considered not justifiable in patients with advanced disease (1). Thus, if the study included histology-only diagnosed dogs a large proportion of dogs would be excluded and survival would be biased to dogs with early stage haemangiosarcoma and longer survival times.

The reported effect of surgical removal on haemangiosarcomas on patient survival time varies widely. Whilst surgical removal has been shown to be potentially curative for cutaneous masses (14), it has been shown to lead to both increase survival time in other locations with varying magnitude (1,7,19,31). Therefore, the effect of surgery on survival time was a variable of interest for the current study. Unfortunately, due to the first opinion nature of these clinics and the fact that many haemangiosarcoma surgeries are highly complex and potentially require specialist surgeons (e.g. liver lobectomy) the effect of specific surgical procedures on survival time could not be assessed. As surgical procedures other than splenectomy were performed infrequently in the current caseload, all surgeries were all grouped into a binary variable which led to a loss of resolution between individual surgical types.

Whilst the mechanism of death was recorded in most EHRs in this study, it was still not possible to disentangle the multiple potential reasons for euthanasia such as perceptions around poor prognosis, financial considerations or whether the perceived acute quality of life deterioration prompted euthanasia. Clinical notes pertaining to euthanasia consults are often brief and not able to capture non-clinical aspects. Therefore, qualitative research might be useful to explore haemangiosarcoma euthanasia considerations.

## Conclusion

This study identified a short survival times for haemangiosarcoma presenting to first opinion practices in the UK. Survival times were shortest for cardiac and hepatic haemangiosarcoma. Tumour location, size, sex and surgical management factors were associated with death with haemangiosarcoma. The wide-ranging survival times seen in haemangiosarcoma cases here (visceral forms ranged from 0-1548 days of survival) indicate that further exploration of tumour biology and potential biomarkers is important, to potentially understand some of the variation between locations and individual dogs. Additionally, the potential effect of perceived prognosis on patient euthanasia should be assessed further.

## Acknowledgements

The authors are grateful to Morris Animal Foundation for their funding of this research. Thanks to Noel Kennedy (RVC) for VetCompass software and programming development. We acknowledge the Medivet Veterinary Partnership, Vets4Pets/Companion Care, Goddard Veterinary Group, CVS Group, IVC Evidensia, Linnaeus Group, Beaumont Sainsbury Animal Hospital, PDSA, Blue Cross, Dogs Trust, Vets Now and the other UK practices who collaborate in VetCompass. We are grateful to The Kennel Club Charitable Trust, Agria Pet Insurance and The Kennel Club for supporting VetCompass.

